# Isolation and characterization of multi-protein complexes enriched in the K-Cl co-transporter 2 from brain plasma membranes

**DOI:** 10.1101/2020.04.30.071076

**Authors:** Joshua L. Smalley, Georgina Kontou, Catherine Choi, Qiu Ren, David Albrecht, Krithika Abiraman, Miguel A. Rodriguez Santos, Christopher E. Bope, Tarek Z. Deeb, Paul A. Davies, Nicholas J. Brandon, Stephen J. Moss

**Affiliations:** Department of Neuroscience, Tufts University School of Medicine, 136 Harrison Ave, Boston 02111; Neuroscience, IMED Biotech Unit, AstraZeneca, Boston, MA 02451; Department of Neuroscience, Physiology, and Pharmacology, University College, London, WC1E, 6BT, UK

**Keywords:** Kcc2, proteome, interactome, phosphorylation

## Abstract

Kcc2 plays a critical role in determining the efficacy of synaptic inhibition, however, the cellular mechanism neurons use to regulate its membrane trafficking, stability and activity are ill-defined. To address these issues, we used affinity purification to isolate stable multi-protein complexes of Kcc2 from the plasma membrane of murine forebrain. We resolved these using Blue-native polyacrylamide gel electrophoresis (BN-PAGE) coupled to LC-MS/MS. Purified Kcc2 migrated as distinct molecular species of 300, 600 and 800 kDa following BN-PAGE. In excess of 90% coverage of the soluble N and C-termini of Kcc2 was obtained. The 300kDa species largely contained Kcc2, which is consistent with a dimeric quaternary structure for this transporter. Intriguingly, lower levels of Kcc1 were also found in this species suggesting the existence of “mixed” Kcc2/Kcc1 heterodimers. The 600 and 800 kDa species represented stable multi-protein complexes of Kcc2. We identified a set of novel structural, ion transporting and signaling protein interactors, that are present at both excitatory and inhibitory synapses, consistent with the proposed association of Kcc2. These included spectrins, ankyrins, and the IP3 receptor. We also identified interactors more directly associated with phosphorylation; Akap5 and Lmtk3. Finally, we used LC-MS/MS on highly purified endogenous plasma membrane Kcc2 to detect phosphorylation sites. We detected 11 sites with high confidence, including known and novel sites. Collectively our experiments demonstrate that Kcc2 is associated with components of the neuronal cytoskeleton and signaling molecules that may act to regulate transporter membrane trafficking, stability, and activity.

## INTRODUCTION

The K^+^/Cl^−^ co-transporter Kcc2 (encoded by the gene SLC12A5) is the principal Cl^-^-extrusion mechanism employed by mature neurons in the CNS (1). Its activity is a pre-requisite for the efficacy of fast synaptic inhibition mediated by Glycine (GLYR) and type A γ-aminobutyric acid receptors (GABA_A_R), which are Cl^−^ permeable ligand-gated ion channels. At prenatal and early postnatal stages in rodents, neurons have elevated intracellular Cl^−^ levels resulting in depolarizing GABA_A_-mediated currents (2). The postnatal development of canonical hyperpolarizing GABA_A_R currents is a reflection of the progressive decrease of intraneuronal Cl^−^ levels that is caused by the upregulation of Kcc2 expression and subsequent activity (3–6). These changes in neuronal Cl^−^ extrusion reflect a sustained increase in the expression levels of Kcc2 after birth, the mRNA levels of which do not reach their maximal levels in humans until 20-25 years of age (7). In addition to this, the appropriate developmental appearance of hyperpolarizing GABA_A_R currents is also in part determined by the phosphorylation status of Kcc2, a process that facilitates its membrane trafficking and activity (8–13).

In keeping with its essential role in determining the efficacy of synaptic inhibition, humans with mutations in Kcc2 develop severe epilepsy soon after birth (14–16). Deficits in Kcc2 activity are also believed to contribute to the development of temporal lobe epilepsy (17, 18), in addition to other traumas including ischemia and neuropathic pain (19, 20). Given the critical role that Kcc2 plays in determining the maturation of inhibitory neurotransmission, subtle changes in its function are also strongly implicated in autism spectrum disorders (ASD) (21), Down syndrome (22), fragile X syndrome (23), and Rett syndrome (24–26).

Here, we isolated and resolved stable protein complexes that contain Kcc2 from highly purified plasma membranes from mouse forebrain, and characterized their composition using an unbiased proteomic approach. We subsequently identified several novel Kcc2-associated proteins. We used the same purification strategy and proteomics to measure Kcc2 phosphorylation. In this way we have developed new insights into the potential mechanisms of Kcc2 stabilization and regulation in the plasma membrane along with measuring the phosphorylation status of Kcc2 on the cell surface.

## Results

### Isolating native stable protein complexes containing Kcc2 from brain plasma membranes

In order to define which proteins are associated with Kcc2 in the neuronal plasma membrane we developed a novel methodology for isolating stable multiprotein complexes enriched in this transporter. To do so, fresh isolated mouse forebrain was homogenized in detergent-free conditions to preserve organelle structure. The resulting homogenate was then subjected to differential gradient-based centrifugation and the plasma membrane fraction was isolated (Figure 1a). To confirm effective fractionation, we immunoblotted for established protein markers for each specific organelle (Figure 1b); HSP90 (cytosol), HSP60 (mitochondria), calreticulin (ER), and n-cadherin (n-Cadh) (plasma membrane). Each organelle fraction showed enrichment for its respective marker protein. Importantly, the plasma membrane fraction was both enriched for n-Cadh and depleted for HSP60 compared to the total lysate, demonstrating efficient separation of plasma membrane and mitochondria, a common contaminant of membranous fractions. Kcc2 was enriched in the ER fraction and highly enriched in the mitochondrial and plasma membrane fractions. The enrichment of Kcc2 in our target fraction - the plasma membrane fraction, was highly encouraging for downstream purification. The presence of Kcc2 in the mitochondrial fraction could be an interesting future area of research, as chloride homeostasis is fundamental for correct mitochondrial function.

**Figure 1.**
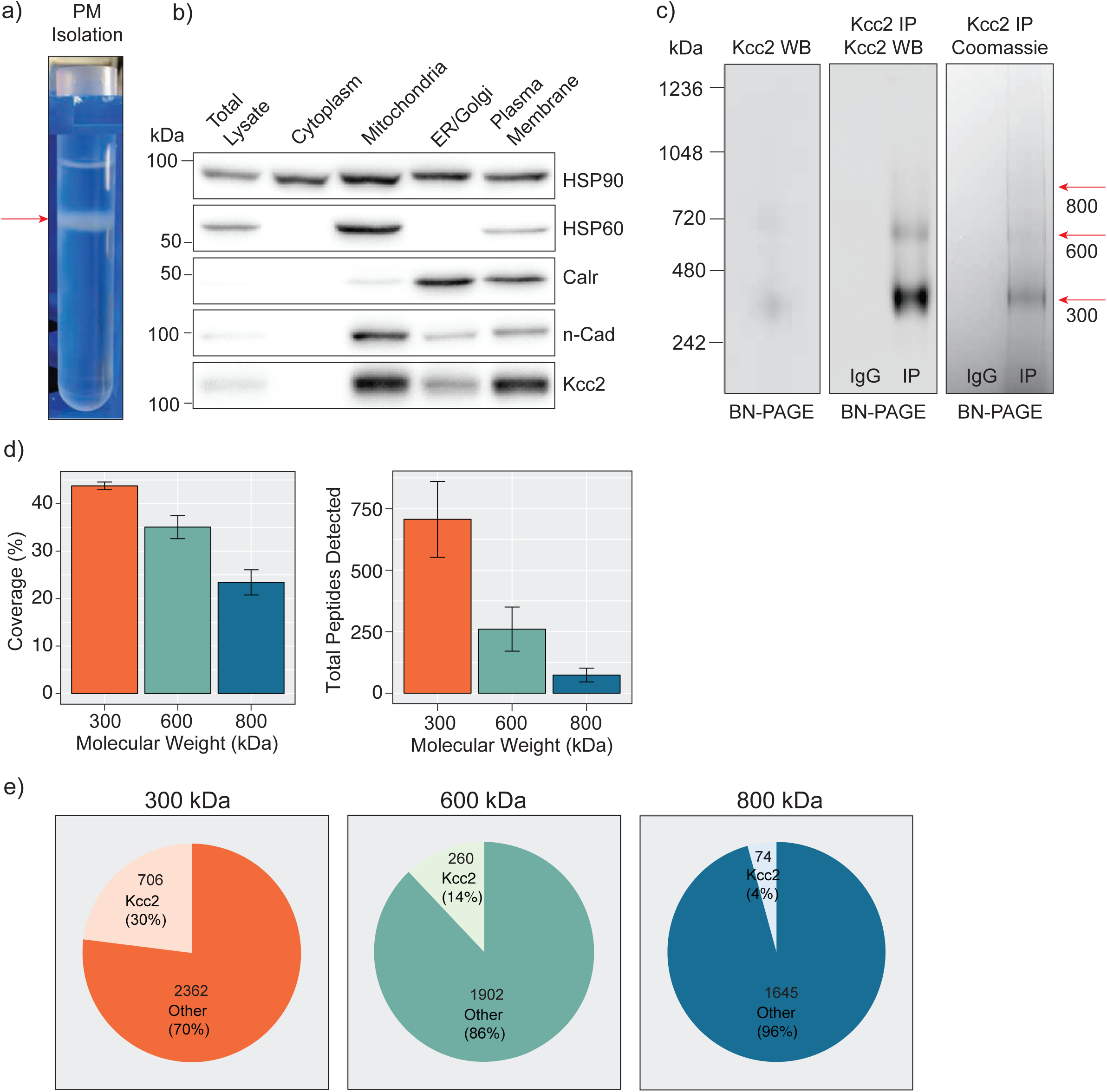
Highly enriched Kcc2-containing protein complexes were isolated from purified plasma membranes from mouse forebrain. **A**. Differential centrifugation of homogenized forebrain from 8-12-week-old mice was used to fractionate cellular organelle and obtain an enriched plasma membrane fraction, visualized here at the final purification stage on a discontinuous sucrose gradient. **B**. Western blots of the organelle fractions were used to measure the abundance of organelle markers; HSP90 (cytosol), HSP60 (mitochondria), calreticulin (ER/Golgi), n-Cadherin (n-Cadh) (plasma membrane). Kcc2 abundance was also measured to confirm Kcc2 enrichment in the plasma membrane fraction. **C**. Blue Native PAGE (BN-PAGE) was carried out on plasma membrane lysates, and immunoprecipitated Kcc2 to resolve the native protein complexes that contain Kcc2. These were correlated with protein bands observed by Coomassie staining. Coomassie-stained bands were excised and used for proteomic analyses. **D.** Detected peptides were mapped to the Kcc2 reference sequence to determine Kcc2 sequence coverage obtained from LC-MS/MS of BN-PAGE protein bands produced following Kcc2 IP. Bar charts of the Kcc2 sequence coverage expressed as a percentage of the full sequence and by total Kcc2 peptides detected (n=4). **E.** Pie charts showing the average number of total Kcc2 peptides detected by LC-MS/MS relative to the average number of total peptides detected for all other proteins in each molecular weight complex.

Following detergent solubilization, membrane fractions were subjected to immunoprecipitation with a monoclonal antibody directed against the intracellular C-terminus of Kcc2 covalently coupled to protein G-coated ferric beads. Initial optimization experiments revealed that we were able to isolate approximately 65% of the Kcc2 available in the lysate and the lysate preparation produced minimal Kcc2 oligomerization when resolved by SDS-PAGE (Figure S1a, b). Purified material was eluted under native conditions using a ‘soft elution buffer’ containing 2% Tween-20 (27), and subjected to Blue-Native-PAGE (BN-PAGE; Figure 1c). We observed well-resolved bands by colloidal Coomassie staining at 300, 600, and 800 kDa. These bands were not seen in control experiments performed with non-immune mouse IgG. An immunoblot of the same sample confirmed that these bands contained Kcc2. Immunoblots of plasma membrane lysates separated by BN-PAGE showed that these distinct Kcc2 complex bands exist in the lysate and are not an artifact of or degraded by purification. Despite the high stringency of our method (with high speed centrifugation and low concentration SDS exposure), no monomeric Kcc2 was observed (∼150 kDa), indicating that Kcc2 was maintained in higher order complexes. This may disrupt low affinity, or transient protein interactions and favor the detection of higher affinity binding partners.

After we confirmed that these well resolved bands contained Kcc2, we assessed their Kcc2 content by LC-MS/MS following trypsin digestion (Figure 1d). The 300, 600, and 800 kDa bands contained an average of 706, 260, and 74 total peptides for Kcc2 respectively, which equated to average coverage of 44%, 35%, and 23% of the total Kcc2 protein sequence respectively. In all three bands, the majority of peptides corresponded to the cytoplasmic and extracellular domains of Kcc2, while peptides corresponding to the transmembrane domains were rarely detected. The failure to detect these regions is consistent with their high hydrophobicity and subsequent low recovery by LC-MS/MS (Figure S2a). We also noted that the 300, 600 and 800 kDa bands contained peptides derived from Kcc1, which were consistently at approximately 6-fold lower levels than those derived from Kcc2. To assess the complexity of each band we compared the average total peptide counts for Kcc2, with the average total peptide counts for all proteins that were detected in our LC-MS/MS spectra (Figure 1e). In the 300, 600, and 800 kDa bands, Kcc2 was the most abundant protein detected, however the proportion of peptides for Kcc2 relative to peptides for all other detected proteins decreased with increasing complex molecular mass, this is likely due to the increased protein complexity of the high molecular weight complexes. Taken together, these results demonstrate our experimental methodology facilitates the isolation of stable high molecular mass protein complexes enriched in Kcc2.

### Analyzing the composition of stable protein complexes of Kcc2

Having confirmed the veracity of our Kcc2 purifications we investigated the reproducibility of the proteins associated with Kcc2 in each of the bands. First, we compiled a data matrix of all the proteins and total peptide counts detected by proteomic analysis in each of our samples (Table S1). We then removed all proteins that were not at least 2-fold enriched in the IP compared to proteins that associated with IgG alone (Table S3) and removed all proteins with fewer than 2 peptide counts in each individual proteomic run. This process is illustrated in Figure 2a.

**Figure 2.**
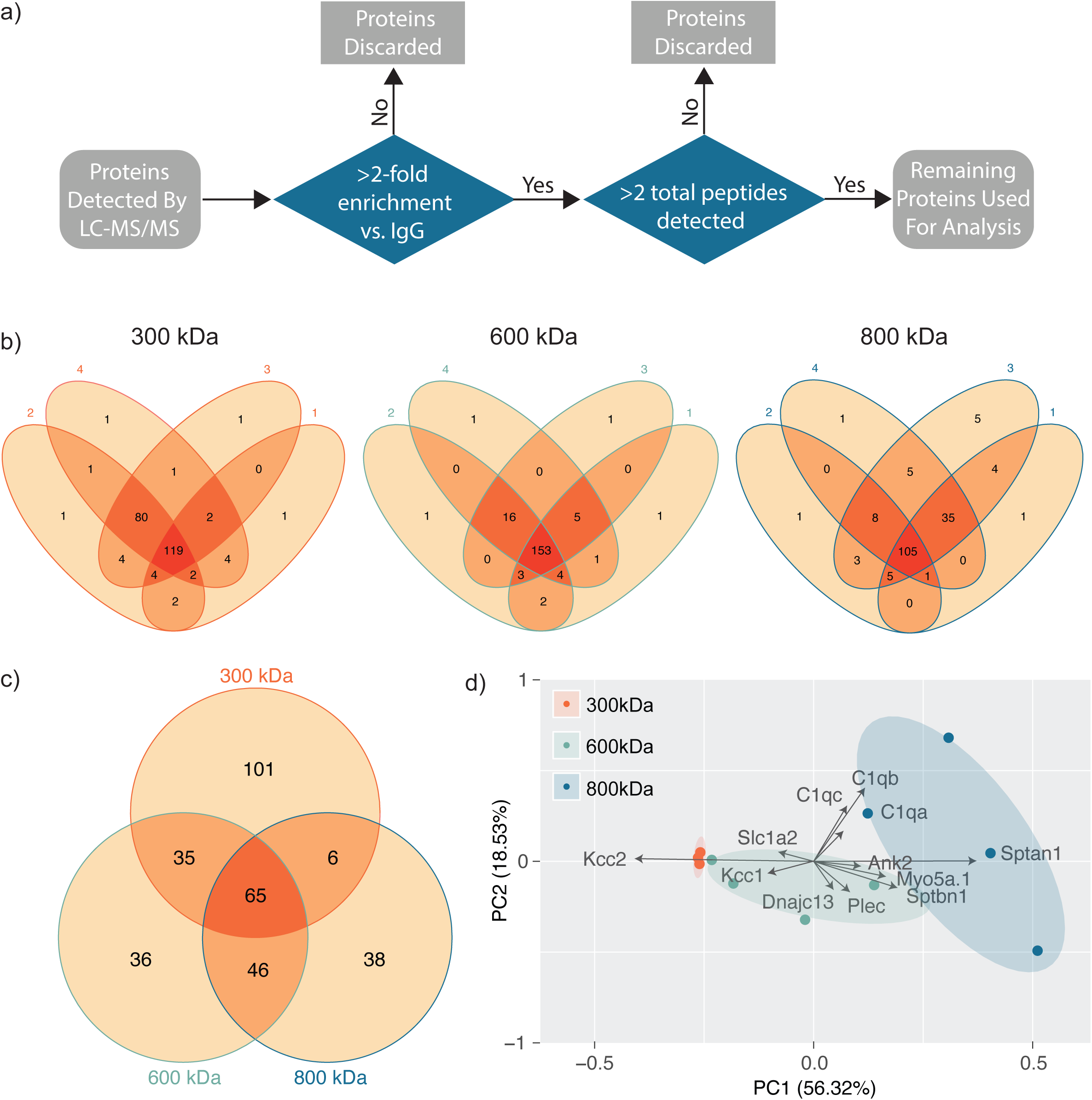
Stable protein complexes contain large amounts of Kcc2 and a robust set of associated proteins, which differ subtly between the different molecular weight bands. **A**. Flow diagram illustrating the first-pass protein filtering process applied prior to quality control analyses. **B**. Venn diagrams showing the number and overlap of Kcc2-associated proteins identified in each of 4 biological replicates in each molecular weight complex. **C**. Venn diagram showing the overlap in associated proteins detected in at least 3 of the 4 replicates for each molecular weight complex. **D**. PCA analysis of each biological replicate for each molecular weight complex based on their protein composition. PCA loadings were also included to show the contribution of each protein to the position of sample on the PCA plot.

Next, we evaluated the reproducibility of the detected proteins in each of the 4 biological repeats for each band (Figure 2b). The vast majority of proteins detected were present in at least 3 of the 4 biological replicates, illustrating the reproducibility of our purifications and analyses. Only proteins detected in at least 3 repeats were taken forward for further analysis as ‘robust binding proteins.’ We then compared these ‘robust binding proteins’ between the 300, 600, and 800 kDa complexes (Figure 2c). There was a high degree of overlap between the 600 and 800 kDa complexes. The 300 kDa complex had considerable overlap with the 600 and 800 kDa complexes, but it also contained many unique proteins. This unique protein pool is likely to be large proteins that have dissociated from Kcc2 complexes.

Finally, we used principle component analysis (PCA) to assess the degree of similarity in binding protein patterns between all 12 of the proteomic samples (Figure 2d). PCA identifies variance between samples and expresses the degree of similarity by proximity in a PCA plot. We did this to both assess the reproducibility of the biological replicates and the degree of difference in the binding proteins for each complex. The ‘robust binding proteins’ were assembled into a matrix of all repeats and detected proteins along with total peptide counts for each. The data was normalized by z-transformation (Table S2) and the PCA plot created using the default settings in the ggfortify package (accessed January 2019) in R (28). The datasets clearly separated into biological repeats of the different mass bands. PC1, which accounted for 56.32% of the variance, was highly positively correlated with amount of Sptan1 and Sptbn1, and negatively correlated with the amount of Kcc2. This is consistent with the detection of more spectrin peptides in the higher molecular weight bands, and a concomitant decrease in the amount of detected Kcc2 due to higher sample complexity. PC2, which accounted for 18.53% of the variance was positively correlated with the detection of more C1qa/b/c peptides. This demonstrates that we have identified a reproducible set of binding proteins for the 300, 600, and 800 kDa bands. While there is a substantial degree of overlap between the interacting proteins from each molecular mass band (Figure 2c), their protein content is different enough that they can be clearly differentiated by PCA.

Next, we performed network analysis on the ‘robust binding proteins.’ We omitted proteins whose peptide counts were lower than 10% of those detected for Kcc2, to remove ‘trace’ binding proteins. The resulting protein lists are shown in Table 1. Network analysis was carried out using data from StringDB for known experimental interactions (29) as previously described (30). We also carried out Gene Ontology (GO) analysis and overlayed the most overrepresented biological process terms (29). We also scaled the node size representing each protein to the total number of peptides detected for each. In this way we were able to visualize Kcc2 networks and subnetworks along with developing insights into the functional groups of proteins that associate with Kcc2 (Figure 3a, b, and c). In the 300 kDa band, a high percentage of the detected peptides were for Kcc2 (an average of 706 across all repeats). Kcc1 (137 peptides) and Cntn1 (80 peptides) were the only proteins detected over our set thresholds. The 600 kDa protein complex contained high levels of Kcc2 (260 peptides), along with Kcc1 (51 peptides). A subnetwork of Sptan1 and Sptbn1 (37 and 34 peptides respectively) also emerged. The associated proteins were largely structural or involved in ion transport. The 800 kDa complex contained high amounts of Kcc2 (74 peptides) along with 3 prominent subnetworks. The most prominent of which contained Sptan1 (65 peptides), Sptbn1 (36 peptides), Sptbn2 (15 peptides), Ank2 (8 peptides), Itpr1 (7 peptides), Acta2 (8 peptides), and Myh10 (10 peptides). There was a subnetwork containing Grm2/3/5 (9, 11, and 15 peptides respectively), and Gabbr2 (8 Peptides). There was also a subnetwork containing C1qa/b/c (23, 27, and 19 peptides respectively). The 800 kDa complex contained more functionally diverse proteins involved in structure (Sptan1, Sptbn1, Sptbn2, Ank2, Acta2, Myh10, Myo5a), ion transport (Slc1a2, Cacna1e, Scn2a, Slc4a4, Slc4a10, and Itpr1), lipid binding (C1qa/b/c, Pitpnm2, Plppr4, and Ralgapa1), and signaling (Grm2/3/5, Gabbr2, Gpr158, Lmtk3, and Akap5).

**Table 1.** Robust binding proteins used for network analysis. Proteins detected by LC-MS/MS were refined by removing IgG associated proteins and removing proteins with fewer than 10% of the peptides detected for KCC2. The table shows the official gene name, Uniprot ID, basic annotation and total peptide counts for the bands at **A.** 300kDa, **B**. 600kDa, **C**. 800kDa. (n=4).

**Figure 3.**
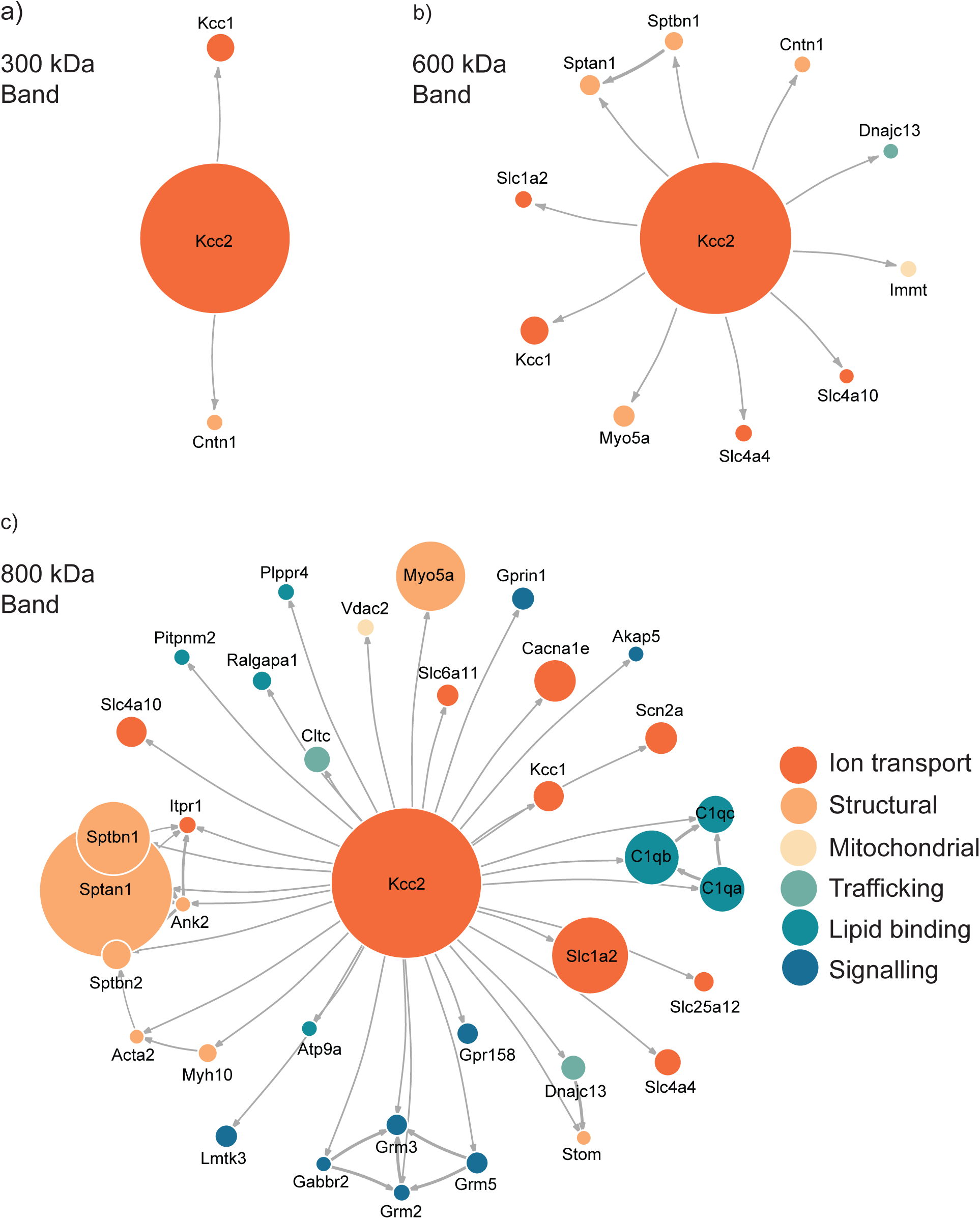
High molecular weight complexes of Kcc2 contain a robust set of proteins from multiple functional classes, with some highly interconnected subnetworks. Protein network diagrams for **A**. the 300 kDa band **B**. the 600 kDa band, and **C**. the 800 kDa band. Only proteins with >10% of the peptides detected for Kcc2 were included to remove ‘trace’ binding proteins. Known interactions were obtained using stringent high confidence, direct experimental association parameters from StringDB. These were used to construct a network diagram of protein nodes and arrows to indicate known interactions. The interactions for each protein with Kcc2 were included as discovered here. The node diameter was scaled relative to the number of peptides detected for each protein. An overlay of gene ontology (GO) terms was used to provide protein classification information (n=4).

### Confirming the association of ‘robust binding proteins’ with Kcc2

Next, we set out to confirm the association of a subset of the newly identified interactors with Kcc2. We chose to focus on the spectrin/ankyrin complex as it was highly prominent and contained both structural and regulatory proteins. We immunoblotted isolated Kcc2 protein complexes resolved by BN-PAGE for Sptan1, Sptbn2, Ank2, and Itpr1 (Figure 4). We also included Cntn1 and Scn2a as specificity controls as they were particularly abundant in the low (300 kDa) and high (800 kDa) bands respectively according to the proteomic analyses. No immunoreactivity was observed in the IgG control lanes. Mirroring the proteomic data Cntn1 showed specificity for the 300 kDa band, whereas Scn2a showed strong specificity for the 800 kDa band. In agreement with the proteomic data, none of the components of the spectrin/ankyrin subnetwork were present in the 300 kDa band. Both Sptan1 and Ank2 were present in the 600 kDa band. Sptan1 was also present in the 800 kDa band along with Sptbn1 and Itpr1.

**Figure 4.**
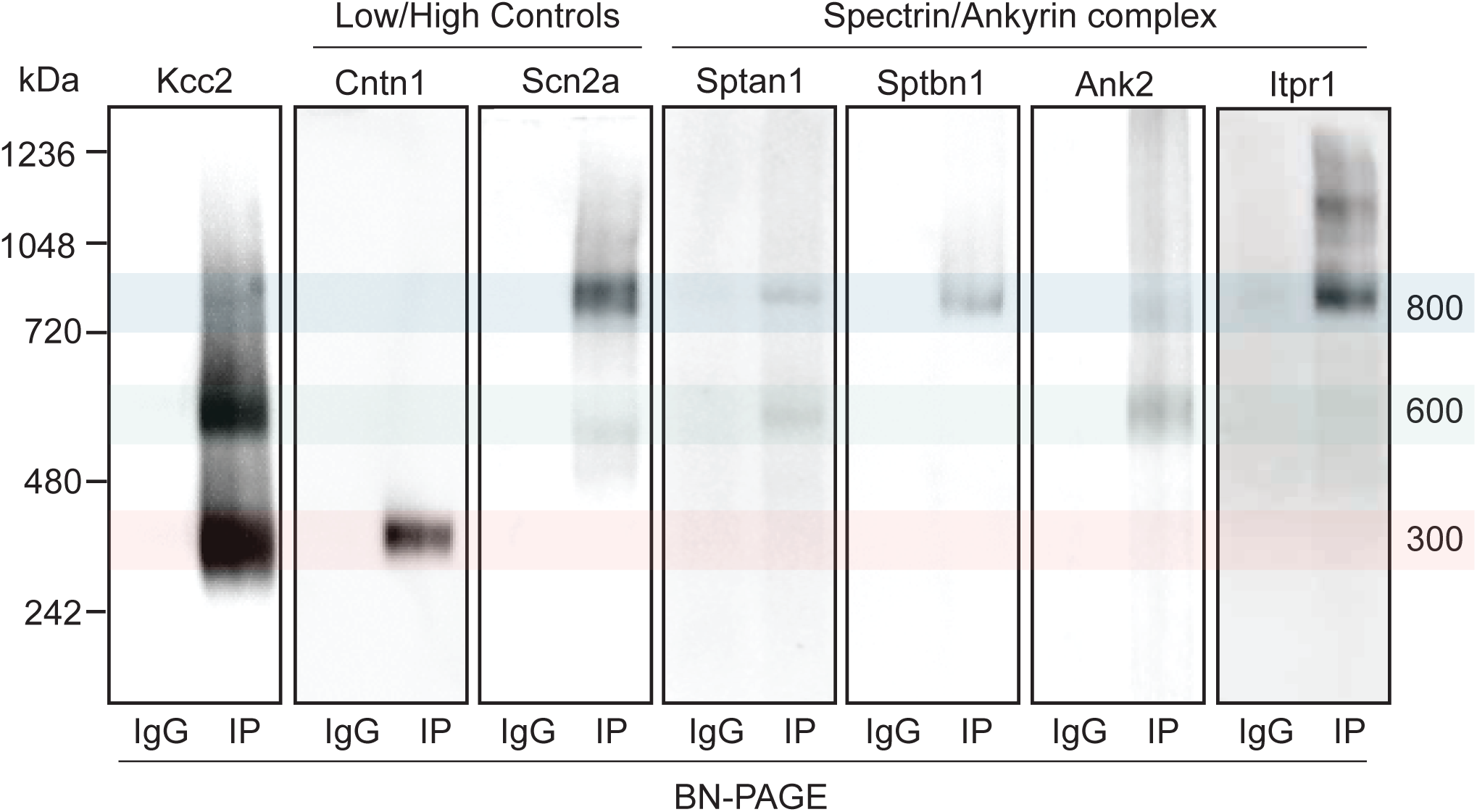
Immunoblots confirm the presence of several interactors in protein complexes that contain Kcc2. **A.** Kcc2 protein complexes were isolated from forebrain plasma membrane fractions of 8-12-week-old mice by immunoprecipitation, resolved by BN-PAGE and immunoblotted for components of the ankyrin/spectrin subnetwork. IPs were loaded adjacent to parallel IgG reactions to demonstrate specificity. Cntn1 and Scn2a were included as a further demonstration of band specificity, as proteomic analysis suggested specific enrichment in the 300 kDa and 800 kDa bands respectively. Transparent overlays mark the location of each Kcc2 complex.

### Kcc2 co-localizes with the spectrin/ankyrin complex on, or close to the neuronal plasma membrane

Having established that Kcc2 forms protein complexes with components of the spectrin/ankyrin complex, we used immunocytochemistry to investigate the proximity of each to Kcc2. We used DIV21 mouse primary cultured cortical/hippocampal neurons infected with CamkII-AAV-GFP to ensure we imaged excitatory neurons and to visualize the morphology of the cell and identify whether the interaction between Kcc2 and the spectrin/ankyrin complex was restricted to specific cellular compartments (Figure 5). Sptan1 exhibited widespread staining along dendrites, which colocalized with puncta of Kcc2. Itpr1 exhibited punctate staining with convincing colocalization with Kcc2 in the dendritic compartment. Ank2 and Sptbn1 were present in dendritic clusters with Kcc2 mostly located on the edges. These results were also reflected in the density plots. Taken together, these data demonstrate that proteins associated in high molecular weight complexes with Kcc2 are also highly colocalized with Kcc2 in neurons.

**Figure 5.**
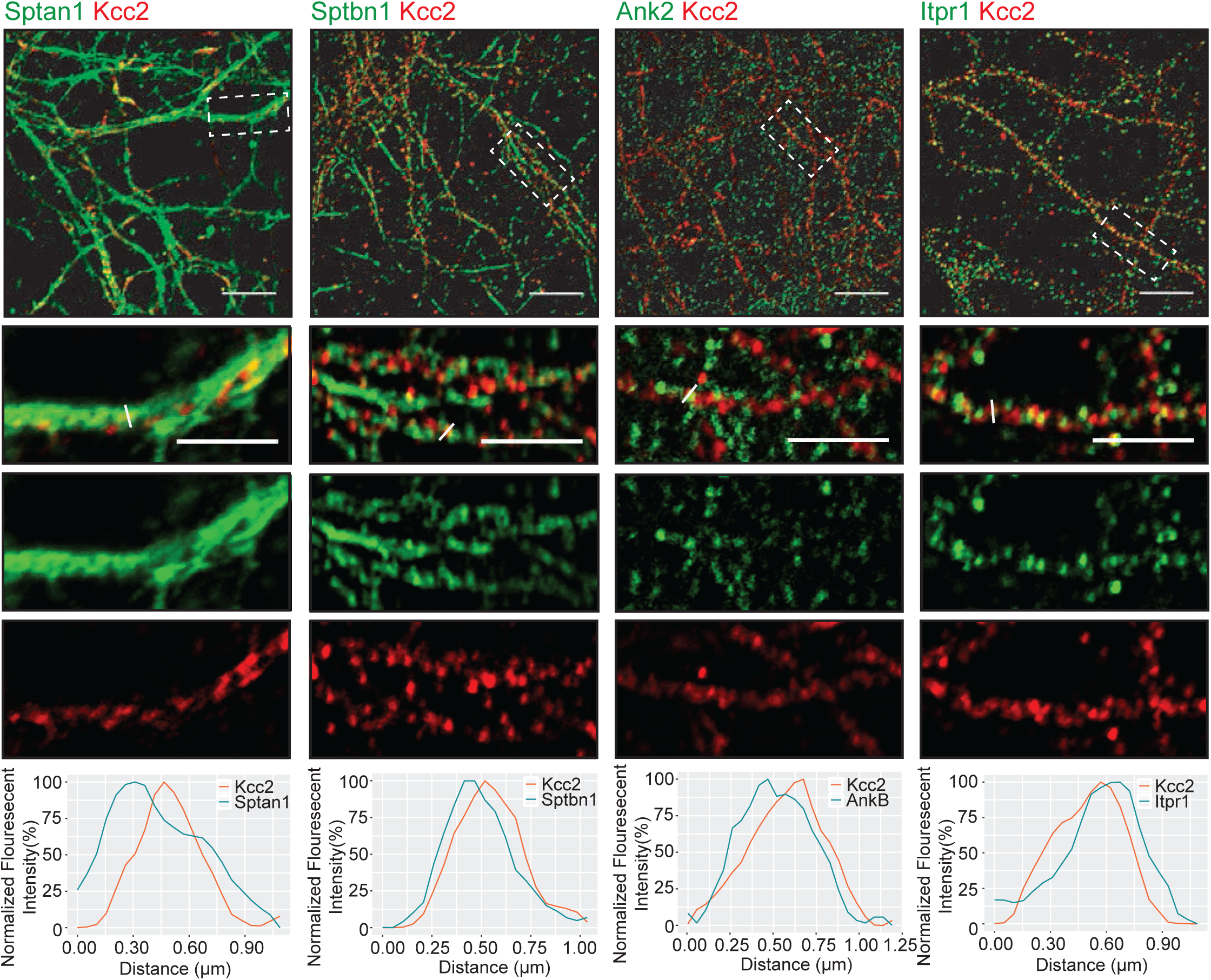
Components of the spectrin/ankyrin subnetwork colocalize with Kcc2 in several neuronal cellular compartments. **A.** Primary cultured neurons from P1 pups were infected with CamKII AAV-GFP at DIV 3 (to visualize cell morphology and to identify excitatory neurons) and fixed at DIV 21. The cells were immunostained for Kcc2 and components of the spectrin/ankyrin subnetwork; Sptan1, Sptbn1, Ank2, Itpr1. Colocalization was measured using line scans (white lines), and represent the normalised fluorescent intensity of single pixels (approximately 1 μm) on dendrites (n=3).

### Kcc2 is highly phosphorylated at multiple sites in the murine plasma membrane

Kcc2 expression levels, membrane trafficking and activity have been shown to be subject to modulation via multiple phosphorylation sites within the C-terminal cytoplasmic domain. As we had succeeded in obtaining highly purified Kcc2 from murine brain plasma membranes, we also analyzed Kcc2 using phosphoproteomics. To do so, we immunoprecipitated Kcc2 from plasma membrane fractions prepared from WT mice as outlined in Figure 1. To limit dephosphorylation purified material was resolved by SDS-PAGE and visualized using Coomassie (Figure 6a). Major bands of 125 kDa were seen with material purified on Kcc2 antibody, but not control IgG. The band was excised, proteins were digested with trypsin, and analyzed by LC-MS/MS. LC-MS/MS confirmed the 125 kDa band was highly enriched for Kcc2 with an average of 884 peptides detected (n=4), which equates to coverage of 53% (Figure S2).

**Figure 6.**
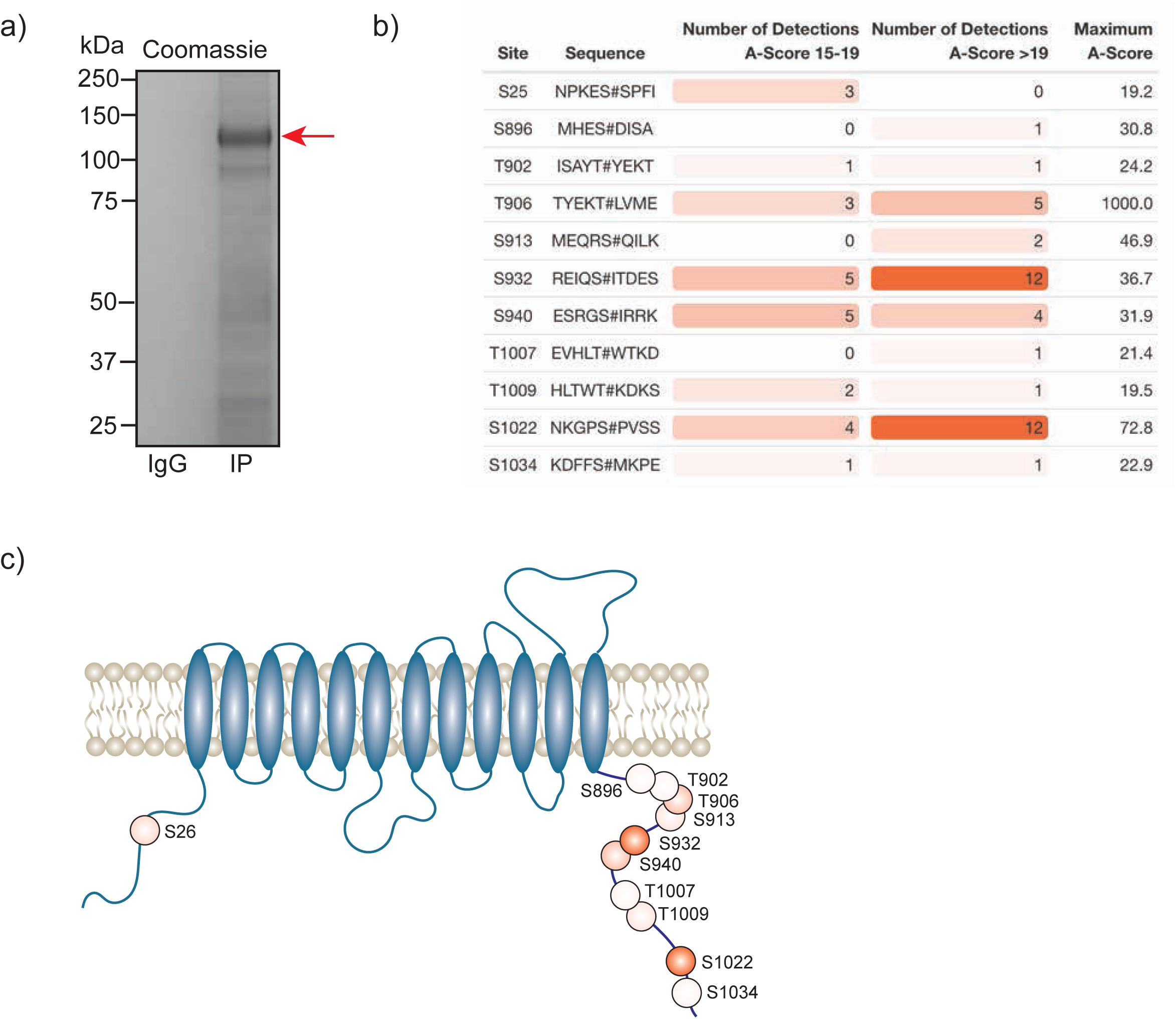
In the neuronal plasma membrane, Kcc2 can be phosphorylated at 11 distinct sites. **A.** Kcc2 was isolated from plasma membrane fractions of mouse forebrain, resolved by SDS-PAGE and visualized using colloidal Coomassie. **B.** Table of detected high confidence Kcc2 phosphorylation sites. Site numbering is according to the mature Uniprot Kcc2 protein sequence. A scores 15-19 and >19 represent a >90% and >99% confidence score respectively. The number of times a particular phospho-peptide was detected at either A-score range was recorded across 4 individual experiments, along with the maximum recorded A-score (n=4). Table cells are color coded according to the number of detections. **C.** Illustrative diagram of high confidence Kcc2 phosphorylation site positions color coded according to the number of detections at an A-score >19. Transmembrane domain positions according to the Uniprot database.

We identified 11 high confidence Kcc2 phosphorylation sites based on A-scores either 15-19 or greater than 19, which equates to 90% or 99% confidence in assignment respectively (Figure 6b). One site (S25/26), is located on the N-terminus of Kcc2, whereas the other 10 sites are located on the C-terminus (S896, T902, T906, S913, T932, S940, T1007, T1009, S1022, and S1034) (Figure 6c). The number of times a phosphopetide was detected with an A score 15-19 or >19 was recorded across 4 biological replicates, to provide a measure of abundance and confidence in each phosphorylation site. We detected the three well characterized sites; T906, S940 and T1007, however T1007 was only detected once with a qualifying A score. The two most regularly detected high confidence phosphosites were S932, and S1022 (detected 18 and 16 times respectively), which are relatively unstudied phosphosites. T906 and S940 were also detected with high confidence several times. Collectively, these results demonstrate that plasma membrane Kcc2 is phosphorylated at multiple sites at the N- and C-termini, at both known and novel sites.

## Discussion

### Isolation of stable multi-protein complexes enriched in Kcc2

The K^+^/Cl^-^ co-transporter, Kcc2, is expressed at the cell surface of mature neurons where it is of fundamental importance for maintaining intracellular chloride levels, and thereby the efficacy of neuronal inhibition. To gain insights into how neurons regulate the plasma membrane accumulation and activity of Kcc2, we developed a method for the isolation of native multiprotein complexes enriched in this transporter from brain plasma membranes. Following plasma membrane isolation by differential centrifugation, immuno-affinity purification, BN-PAGE/SDS-PAGE and LC-MS/MS we were able to isolate highly purified preparations of Kcc2. When resolved by SDS-PAGE we obtained a single band at approximately 140 kDa. When resolved by BN-PAGE we observed major molecular mass species of 300, 600, and 800 kDa. This methodology provided very high sequence coverage of Kcc2 (approximately 45% of the total sequence, and 90% of the intra- and extracellular soluble regions), maximizing the probability of detecting associated proteins and post-translational modifications. It could therefore prove to be a beneficial methodology for the isolation of other plasma membrane proteins from brain tissue, that not only maintains protein complexes, but also post-translational modifications.

### Kcc2 exists as dimers in the neuronal plasma membrane

Kcc2 consists of 1139 amino acids including the signal sequence, which equates to a mass of 126 kDa without post-translational modifications. Kcc2 is, however, subject to extensive post-translational modification including glycosylation at six sites so is often observed in a band of around 140 kDa (31). Previous work has shown extensive aggregation of Kcc2 following SDS-PAGE that results in a band at 250 kDa, however this depends on whether it is heterologously expressed or the manner in which it is prepared from neuronal tissues (32). Indeed, we see no aggregation in our samples following SDS-PAGE, possibly due to the use of low detergent concentrations and samples immediately processed for SDS-PAGE without freeze thaw cycles. Interestingly, when resolved by BN-PAGE, Kcc2 protein complexes form distinct bands at 300 kDa, 600 kDa and 800 kDa. The 300 kDa band contains approximately 6-fold more peptides for Kcc2 than the next most abundant protein, indicating that this band consists largely of Kcc2. At 300 kDa, this would indicate that in the plasma membrane Kcc2 exists as homodimers, which is consistent with recently published studies on recombinant Kcc2 molecules visualized using cryo-EM (31) Interestingly, the next most abundant protein in the 300 kDa species was Kcc1, potentially indicating that Kcc1/Kcc2 heterodimers are present in the brain, albeit at lower levels than Kcc2 homodimers. Kcc1 has also been shown to form dimers by cryo-EM (33) and this is consistent with the notion from studies in oocytes that suggest that Kcc2 can form heterodimers with other Kcc-proteins (34). Whether these potential heterodimers are functional is unclear but is an exciting focus for future research.

### Kcc2 forms stable multi-protein complexes in the plasma membrane

The most abundant protein in the 600 kDa and 800 kDa bands was consistently Kcc2, confirming that these are legitimate Kcc2-containing protein complexes. Significantly the ratio of the associated proteins relative to Kcc2 increased with molecular mass. Given the stringency of our methodology and the use of native conditions these species represent stable-protein complexes that primarily reside in the plasma membrane. The proteins detected by a previous proteomic analysis of Kcc2 complexes (35) overlapped with 13% and 16% of the proteins we detected in the 600 kDa and 800 kDa bands respectively. This low degree of overlap is likely due to the methodological differences between the studies; the previous work extracted Kcc2 complexes from a crude membrane preparation and used on-bead digest prior to LC-MS/MS, rather than complexes purified from a plasma membrane fraction resolved by BN-PAGE, then distinct bands excised for analysis. The previously presented work is possibly more likely to detect transient interactions due to less sample processing, but is also far less stringent, and results in a highly complex sample being introduced to the LC-MS/MS, which can significantly impact resolution (36, 37). However, the method presented here ensured a sample of reduced complexity analyzed by LC-MS/MS, resulting in a higher resolution snapshot of the proteins associated with high affinity with Kcc2 on the neuronal plasma membrane. Similarly to Mahadevan *et al*., we also did not detect Neto2 (36), Grik2 (Gluk2) (37), Epb41l1 (4.1n) (38), Arhgef7 (Beta-pix) (39), Rcc1 (40) or with the signal transduction molecules; Pkc, Wnk, Spak or Osr (8, 41). This could be either due the transient or low affinity nature of these interactions, or that these interactors are less abundant in complex with Kcc2 than the interactors presented here, as unbiased methodologies provide information on associated proteins in rank order of abundance. We did, however, detect GABA_B_Rs which have previously shown to be associated with Kcc2 (42), further confirming the veracity of our purification strategy.

### Kcc2 is associated with excitatory and inhibitory synapses

Previous work has shown that Kcc2 is located at or in close proximity to excitatory synapses (35, 43, 44). In agreement with this we detected well characterized excitatory synapse proteins such as Cntn1 (45), Grm2/3 (46, 47), Myo5a (48). Interestingly, we also identified several proteins associated with inhibitory synapses including Cacna1e (49), and Cyfip1 (50). Grm5 has been detected in high abundance at both excitatory and inhibitory synapses (46, 49). Taken together, these data suggest that Kcc2 is not located exclusively at either type of synapse but is likely to be positioned at or in close proximity to both. This is consistent with the suggestion that Kcc2 may play a role in the regulation of excitatory and inhibitory synapses (51). It is also of note that many of the proteins we detected are protein products of autism- and epilepsy-risk genes.

### Kcc2 is associated with the spectrin/ankyrin cytoskeleton

The most prominent subnetwork that we identified associated with Kcc2 was the spectrin/ankyrin complex, which consisted of Sptan1, Sptbn1, Sptbn2, Ank2, and Itpr1 (52). Sptan1, the most abundant binding protein in the higher molecular weight complexes, is a structural protein that forms dimers with a β-spectrin isoform such as Sptbn1. These spectrin dimers interact with actin filaments and ankyrins, such as Ank2, to regulate the expression of plasma membrane proteins, including calcium channels (including Cacna1e) (53), sodium channels (including Scn2a) (54), potassium channels, and Itpr1 (55). Cntn1 is also thought to interact with spectrins either directly or via Caspr1 (56). The importance of this complex is reflected in the debilitating diseases associated with spectrin/ankyrin mutations. Sptan1 mutation is associated with epilepsy (57), and Sptbn1 is implicated in ASDs (58). Both Ank2 and Itpr1 are high risk ASD genes identified by SFARI. Sptan1 cleavage by calpains, as also occurs with Kcc2 (59), is an established marker of neuronal damage (60).

Our data suggests that Kcc2 most likely interacts directly with Sptan1 or Ank2, as both are present in high abundance in the 600 kDa band with Kcc2. In contrast, Sptbn1 and Itpr1 are likely to require the binding of Sptan1 or Ank2 to associate with Kcc2, as they are most highly abundant in the 800 kDa band, and absent in the 600 kDa band (Figure 4.) This hypothesis is also consistent with the shift in molecular weight observed in the Kcc2 complex. A Kcc2 dimer of approximately 300 kDa would run at approximately 600 kDa with an associated Sptan1 protein (285 kDa), or Ank2 (120-420 kDa depending on splice variant). The subsequent association of either Sptbn1 (275 kDa), or Itpr1 (310 kDa) would shift the mass to around 800 kDa. Both Sptan1 and Ank2 have a variety of binding domains where interactions with Kcc2 could occur, such as SH3 domains (Sptan1) (61) and the membrane binding domain containing the Ank repeat sequences of Ank2 (62), which has been shown to interact with other ion channels/transporters (63).

The association of Kcc2 with the spectrin/ankyrin complex would not only stabilize Kcc2 in the plasma membrane, but also provide a link to signaling processes via Itpr1. Itpr1 is present at PM/ER junctions and following activation by G-protein coupled receptors (GPCR), releases Ca^+^ from the ER. High Ca^+^ concentration would lead to localized PKC activation, which has been shown to phosphorylate Kcc2 at multiple sites (8).

### Plasma membrane populations of Kcc2 are phosphorylated on up to 11 residues

We used the same purification techniques to analyze Kcc2 phosphorylation from the neuronal plasma membrane. Phosphorylation plays a key role in regulating the transporter activity. We detected 11 residues including the well characterized sites; T906, S940, and T1007. These play a critical role in regulating Kcc2 function (13, 64). Interestingly, T1007 was only detected above the accepted A-score threshold once. T1007 phosphorylation is inversely correlated Kcc2 function (65), and therefore this could suggest that T1007 is largely absent in PM-associated Kcc2, where it is active.

In addition to the well characterized sites we also detected 8 further phosphorylation sites. These included S25/S26 and S1022 consistent with recent publications (31, 66). We also identified 5 further sites of phosphorylation; S896, T902, S913, S932, T1009, and S1034. S932 and S1022 were the most frequently detected phosphosites, indicating that Kcc2 in the PM is highly phosphorylated at these residues. S932 is in close proximity to the well characterized S940 site, which is vital for correct Kcc2 function (67). S1022 lies adjacent to, or within the isotonic domain of Kcc2, suggesting that its phosphorylation may impact its basal activity. (68).

In conclusion we have demonstrated a highly efficient method for isolating Kcc2 from murine plasma membranes, in a highly phosphorylated form within its native protein complex. It confirms Kcc2 is in either homo- or hetero-dimeric conformation with Kcc1. These dimers associate with a spectrin/ankyrin subnetwork that is likely to not only stabilize it in the plasma membrane but provide an environment for regulating its phosphorylation at the 11 potential sites.

### Experimental Procedures

Unless otherwise stated chemicals were obtained from Sigma-Aldrich, St. Louis, MO, USA.

#### Animals

Animal studies were performed according to protocols approved by the Institutional Animal Care and Use Committee of Tufts Medical Center. 8-12-week-old male and female mice were kept on a 12-hour light/dark cycle with *ad libitum* access to food and water.

#### Antibodies

The following antibodies were used for immunoprecipitation (IP), immunoblot (IB), or immunocytochemistry (Icc): Ank2 (mouse, Icc, Invitrogen 33-3700), Ank2 (rabbit, IB, Bioss BS-6967R-TR), Cntn1 (rabbit, IB), Abcam Ab66265), Itpr1 (rabbit, IB/Icc, Alomone Acc-019), Kcc2 (mouse, IP/Icc, Neuromab 75-013), Kcc2 (rabbit, IB/Icc, Millipore 07-432), Scn2a (mouse, Neuromab 75-024), Scn2a (rabbit, IB, Alomone ASC-002), Sptan1 (mouse, Icc, Abcam Ab11755), Sptan1 (rabbit, IB, Cell Signaling 21225), Sptbn1 (rabbit, IB/Icc, Abcam Ab72239), α-Tubulin (mouse, IB, Sigma T9026).

#### Immunoblotting

Sodium dodecyl sulphate polyacrylamide gel electrophoresis (SDS-PAGE) was carried out as previously described (69). Briefly, proteins were isolated in RIPA lysis buffer (50 mM Tris, 150 mM NaCl, 0.1% SDS, 0.5% sodium deoxycholate and 1% Triton X-100, pH 7.4) supplemented with mini cOmplete protease inhibitor and PhosSTOP phosphatase inhibitor tablets. Protein concentration was measured using a Bradford assay (Bio-Rad, Hercules, CA, USA). Samples were diluted in 2x sample buffer and 20 μg of protein was loaded onto a 7-15% polyacrylamide gel depending on the molecular mass of the target protein. After separation by SDS-PAGE, proteins were transferred onto nitrocellulose membrane. Membranes were blocked in 5% milk in tris-buffered saline 0.1% Tween-20 (TBS-T) for 1 hour, washed with TBS-T, and then probed with primary antibodies diluted in TBS-T (dilution and incubation time dependent on the antibody). The membranes were washed and incubated for 1 hour at room temperature with HRP-conjugated secondary antibodies (1:5000 – Jackson ImmunoResearch Laboratories, West Grove, PA, USA). Protein bands were visualized with Pierce ECL (ThermoFisher) and imaged using a ChemiDoc MP (Bio-Rad). Band intensity was compared to α-tubulin as a loading control.

#### BN-PAGE

For blue native poly-acrylamide gel electrophoresis (BN-PAGE), proteins were isolated using Triton lysis buffer (150mM NaCl, 10mM Tris, 0.5% Triton X-100, pH 7.5). Samples were diluted in 4x NativePAGE sample buffer and G250 additive. Samples were loaded onto 4-16% NativePAGE gradient gels and run using a mini gel tank (ThermoFisher, Waltham, MA, USA). Gels were run for approximately 2 hours and then prepared for coomassie staining or immunoblotting. For coomassie staining the gels were fixed in 50% ethanol and 10% acetic acid, washed in 30% ethanol, washed in water, then stained with EZ blue stain. The gels were destained in ultrapure water, imaged using a ChemiDoc MP (Bio-Rad), and bands were excised for liquid chromatography tandem mass spectrometry (LC-MS/MS). For immunoblotting, proteins were transferred to PVDF membrane overnight. The membranes were then fixed in 8% acetic acid, washed with ultrapure water and air-dried before being briefly destained with 100% methanol. The membranes were then blocked, immunoblotted, and imaged as described above.

#### Plasma membrane isolation

Plasma membrane isolation from mouse forebrain was carried out using a modified version of a previously described method (70). Briefly, mouse forebrain was rapidly isolated and placed in ice-cold starting buffer (225mM mannitol, 75mM sucrose, 30mM Tris-HCl (pH 7.4). The tissue was then transferred to ice-cold isolation buffer (225mM mannitol, 75mM sucrose, 0.5% (wt/vol) BSA, mM EGTA, 30mM Tris-HCl, pH 7.4) supplemented with mini cOmplete protease inhibitor and PhosSTOP. 5-7 brains were used per immunoprecipitation. The brains were homogenized in the isolation buffer using 14 strokes of a Dounce homogenizer. The samples were transferred to centrifuge tubes for centrifugation, which was carried out at 4°C throughout. The samples were initially spun at 800xg for 5 minutes to remove nuclei and non lysed cells. The pellet was discarded, and the supernatant was spun again at 800xg for 5 minutes to remove residual nuclei and non lysed cells. The supernatant was transferred to high speed centrifuge tubes and spun at 10000xg for 10 minutes to remove mitochondria. The pellet was discarded, and the supernatant spun again at 10000xg for 10 minutes to remove mitochondrial contamination. The supernatant was then spun at 25000xg for 20 minutes to pellet plasma membranes. The pellet was resuspended in starting buffer and spun again at 25000xg for 20 minutes to remove cytosolic and ER/Golgi contamination. Finally, the plasma membrane fraction was purified on a discontinuous sucrose gradient (53%, 43%, 38% sucrose), and spun at 93000xg for 50 minutes to remove residual contamination from cytosol and other membranous compartments.

#### Protein purification

Protein G Dynabeads (ThermoFisher) were washed 3 times with phosphate buffered saline with 0.05% Tween-20 (PBS-Tween). The beads were resuspended in PBS-Tween and incubated overnight at 4°C with antibodies for the target protein at an experimentally predetermined bead:antibody ratio (Figure S1). The antibody was crosslinked onto the beads by washing twice with 0.2M triethanolamine (pH 8.2) (TEA), and then incubated for 30 minutes with 40mM dimethyl pimelimidate (DMP) in TEA at room temperature. The beads were transferred to 50mM Tris (pH7.5) and incubated at room temperature for 15 minutes. The beads were then washed 3 times with PBS-Tween and resuspended in solubilized plasma membranes in ice cold Triton lysis buffer, supplemented with mini cOmplete protease inhibitor and PhosSTOP. The immunoprecipitation reaction was incubated overnight at 4°C. The beads were then washed 3 times with PBS-Tween and eluted either with 2x sample buffer (for SDS-PAGE) or soft elution buffer (0.2% (w/V) SDS, 0.1% (V/V) Tween-20, 50 mM Tris-HCl, pH = 8.0) (27) (for BN-PAGE).

#### Protein and phosphosite identification

Excised gel bands were cut into 1 mm^3^ pieces and subjected to modified in-gel trypsin digestion, as previously described (71). Briefly, gel pieces were washed and dehydrated with acetonitrile for 10 minutes and then completely dried in a speed-vac. Gel pieces were rehydrated with 50 mM ammonium bicarbonate solution containing 12.5 ng/µl modified sequencing-grade trypsin (Promega, Madison, WI, USA) and incubated for 45 minutes at 4°C. The excess trypsin solution was removed and replaced with 50 mM ammonium bicarbonate solution. Samples were then incubated at 37°C overnight. Peptides were extracted by washing with 50% acetonitrile and 1% formic acid. The extracts were then dried in a speed-vac (∼1 hour). The samples were stored at 4°C until analysis. Before analysis the samples were reconstituted in 5 - 10 µl of HPLC solvent A (2.5% acetonitrile, 0.1% formic acid). A nano-scale reverse-phase HPLC capillary column was created by packing 2.6 µm C18 spherical silica beads into a fused silica capillary (100 µm inner diameter x ∼30 cm length) with a flame-drawn tip (72). After equilibrating the column each sample was loaded via a Famos auto sampler (LC Packings, San Francisco, CA, USA) onto the column. A gradient was formed, and peptides were eluted with increasing concentrations of solvent B (97.5% acetonitrile, 0.1% formic acid). As each peptide was eluted, they were subjected to electrospray ionization and then entered into an LTQ Orbitrap Velos Pro ion-trap mass spectrometer (Thermo Fisher Scientific, San Jose, CA, USA). Eluting peptides were detected, isolated, and fragmented to produce a tandem mass spectrum of specific fragment ions for each peptide. Peptide sequences (and hence protein identity) were determined by matching protein or translated nucleotide databases with the acquired fragmentation pattern using Sequest (ThermoFinnigan, San Jose, CA, USA) (73). For phosphopetide detection the modification of 79.9663 mass units to serine, threonine, and tyrosine was included in the database searches. Phosphorylation assignments were determined by the A score algorithm (74). All databases include a reversed version of all the sequences and the data was filtered to between a one and two percent peptide false discovery rate.

#### Primary neuron culture

Mouse cortical and hippocampal mixed cultures were created from P0 mouse pups as previously described (75). Briefly, P0 mice were anesthetized on ice and the brains removed. The brains were dissected in Hank’s buffered salt solution (Invitrogen/ThermoFisher) with 10 mM HEPES. The cortices and hippocampi were trypsinized and triturated to dissociate the neurons. Cells were counted using a hemocytometer and plated on poly-l-lysine-coated 13mm coverslips in 24-well plate wells at a density of 2 × 10^5^ cells/ml in Neurobasal media (Invitrogen/ThermoFisher). At days *in vitro* (DIV) 18, cells were fixed in 4% paraformaldehyde in PBS for 10 minutes at room temperature. They were then placed in PBS at 4°C until being processed for immunocytochemistry.

#### Immunocytochemistry

Fixed primary neurons were permeabilized for 1 hour in blocking solution (3% BSA, 10% normal goat serum, 0.2 M glycine in PBS with 0.1% Triton-X100). Cells were exposed to primary and then fluorophore-conjugated secondary antibodies diluted in blocking solution for 1h each at room temperature. The coverslips were then washed in PBS, dried, and mounted onto microscope slides with Fluoromount-G (SouthernBiotech, Birmingham, AL, USA). The samples were imaged using a Nikon Eclipse Ti (Nikon Instruments, Melville, NY, USA) or Leica Falcon (Leica Microsystems, Buffalo Grove, IL, USA) confocal microscope using a 60x oil immersion objective lens. Image settings were manually assigned for each fluorescent channel. For image processing, the background was subtracted for each fluorescent channel and the median filter was applied (Radius = 1 pixel) on Fiji Software (76). The line scans (white lines) used for protein localization were generated using the PlotProfile function in FIJI and represent the fluorescent intensity of single pixels against the distance of a manually drawn line (approximately 1 μm) on dendrites.

#### Densitometry

For Western blot analysis, bands from raw images were analyzed using the FIJI densitometry features. Biological replicates were run on the same gels for comparison, and area under the curve was calculated for each band. Average signal and standard error were calculated for each treatment group and ANOVA carried out using R for statistical comparison of protein expression levels.

#### Bioinformatics and Statistics

Detected peptides for each Uniprot ID were compared to the Uniprot mouse reference sequence for annotation using R. Kcc2 peptide sequences were aligned to the mouse Kcc2 reference sequence (Uniprot ID: Q91V14) using the Multiple Sequence Alignment (MSA) package in R (accessed January 10^th^, 2019). Proteins with significant affinity for IgG (less than 2-fold enrichment in the IP vs the IgG control) were removed and consideration given to previous experiments detecting proteins prone to binding to IgG alone (77, 78). For first-pass filtering for quality control analysis, proteins with fewer than 2 total peptides and not detected in at least 3 of the 4 replicates were discarded. Venn diagrams were produced using the Vennerable package in R (accessed January 10^th^, 2019), and only proteins detected in all repeats were considered for downstream analysis. These total peptide counts for the lists of proteins contained within each gel band (Table S1) were normalized by z-transformation (Table S2) and used for Principle Component Analysis (PCA), which was carried out using PCA functions in the ggfortify package in R (accessed January 10^th^, 2019). The protein lists for each band were ordered according to total peptide counts, a measure for relative abundance, proteins with <10-fold fewer peptide counts than those detected for Kcc2 were removed as they were considered ‘trace’ binding partners. The protein lists were compared against the latest version of the StringDB database (29) to establish known interactions and annotations for each protein using only high confidence, experimental evidence. The interaction for each protein with Kcc2 was imputed and network diagrams were constructed in R using the igraph package (accessed February 1^st^, 2019).

### Data Availability Statement

All the data are contained within this manuscript.

## Acknowledgements

S.J.M. is supported by National Institutes of Health (NIH)–National Institute of Neurological Disorders and Stroke Grants NS051195, NS056359, NS081735, R21NS080064 and NS087662; NIH–National Institute of Mental Health Grant MH097446,

## Author Contributions

JLS and SJM conceptualized the project, analyzed the data and wrote the paper. GK performed immunocytochemistry experiments. CC and QR performed western blot experiments. MARS and CEB maintained the mouse colony and performed genotyping. TZD and NJB edited the paper.

